# Genome-wide DNA methylation study reveals specific signatures in the affected arterial tissue of giant cell arteritis patients

**DOI:** 10.1101/2025.02.27.640503

**Authors:** Gonzalo Borrego-Yaniz, Ana Márquez, Elkyn Estupiñán-Moreno, Laura C. Terrón-Camero, Miguel A. González-Gay, Santos Castañeda, Giuliana Guggino, David Saadoun, Pietro Lio, Simona Fontana, Martina Bonacini, Alessandro Rossi, Alberto Cavazza, Francesco Muratore, Carlo Salvarani, Nicolo Pipitone, Javier Martin, Stefania Croci, Lourdes Ortiz-Fernández

## Abstract

**Objectives:** Giant cell arteritis (GCA) is a large-vessel vasculitis, potentially causing complications such as blindness and strokes. This study aims to gain insights into the pathogenesis of GCA by identifying specific DNA methylation signatures in the arterial tissue of patients with this vasculitis.

**Methods:** DNA methylation profiling was analyzed in 79 temporal artery biopsy samples (69 patients with GCA and 10 controls) by performing an epigenome-wide association study (EWAS). Differential analysis was performed to identify differentially methylated positions (DMPs) and regions (DMRs). Lastly, we compared our findings with previous transcriptomics and epigenomics studies on GCA-affected arteries.

**Results:** EWAS identified 3,644 DMPs (FDR < 0.05, |Δβ| > 0.3), indicating a profound alteration within GCA-affected arterial tissue. These DMPs were annotated to 1,517 potentially dysregulated genes. 282 additional genes were identified by annotation of significant DMRs. Pathway enrichment analysis revealed a significant alteration of inflammatory mechanisms, such as interleukins 2 and 7, as well as pathways related to vascular remodeling. Omics study comparison revealed 37 genes consistently affected across datasets, many of them linked to immune signaling and T cell regulation. Notably, markers of exhausted T cells, including *SLAMF6* and *HAVCR2*, were present among them.

**Conclusions:** Our study identified GCA-specific DNA methylation signatures in arterial tissue, revealing disrupted inflammatory and vascular pathways, and suggesting the involvement of exhausted T cells in this condition. These findings offer new insights into GCA pathogenesis and provide new potential targets for the treatment of this debilitating disease.

**Key messages:** *What is already known on this topic:*

□ Affected arteries by giant cell arteritis (GCA) exhibit unique epigenetic signatures, reflecting significant disruptions in gene regulation. However, small sample sizes have constrained the clinical translation and broader interpretation of these findings, leaving key gaps in understanding GCA pathogenesis and identifying therapeutic targets.

*What this study adds:*

□ This study provides the largest-to-date epigenome-wide DNA methylation profiling in GCA-affected arteries, uncovering thousands of epigenetic changes and revealing profound disruptions in inflammatory and vascular pathways, including IL-2, IL-7, and CXCR4 signaling.
□ We compiled 37 genes consistently affected across different omics datasets in GCA-affected arteries, providing a robust list of candidates for further research into GCA pathogenesis and treatment.
□ Our findings provide evidence of T cell exhaustion in GCA-affected arteries, supported by consistent changes in key markers such as *SLAMF6* and *HAVCR2* (TIM-3), suggesting a novel mechanism of immune dysregulation in GCA.

*How this study might affect research, practice or policy:*

□ This study nominates exhausted T cells, the NLRP3 inflammasome, and CXCR4 signaling as novel contributors in GCA pathogenesis, suggesting new therapeutic targets for further research in the treatment of this disease.

## Introduction

Giant cell arteritis (GCA) is the most common form of vasculitis in individuals over 50 years, characterized by chronic inflammation of the aorta and its branches. This condition carries significant morbidity and mortality, with complications such as irreversible vision loss and stroke [1]. Despite its clinical impact, the molecular mechanisms underlying GCA pathogenesis remain insufficiently understood, limiting the development of improved treatments and management strategies [2, 3].

Pathological hallmarks of GCA are the loss of immune privilege in the arterial tissue, sustained inflammatory damage leading to severe vascular remodeling, and the neovascularization of the extracellular matrix [4]. These processes create a chronic inflammatory microenvironment that sustains the malfunctioning immune response in the vascular tissue. Different studies have shown genetic risk factors and reprogrammed gene regulation associated with this condition [5–8], showing a complex contribution of genetic and epigenetic drivers to GCA pathogenesis.

Epigenetics is a dynamic mechanism to regulate gene expression, affecting development, tissue differentiation and cellular responsiveness [9]. DNA methylation is the most studied epigenetic marker, having been associated with pathogenic conditions such as immune-mediated inflammatory diseases (IMIDs) [10]. Epigenetic signatures associated with disease progression hold great potential for the discovery of new therapeutic targets due to their reversible nature and potential to address disease mechanisms influenced by genetics, lifestyle and environmental factors [11]. In GCA, a previous study revealed thousands of methylation changes present in temporal artery biopsies [12], however, due to limited sample size and CpGs coverage, there is an incomplete understanding of the role of this mechanism in GCA pathogenesis.

In this study, we conducted a comprehensive DNA methylation analysis of GCA-affected arteries. By leveraging genome-wide profiling in the largest GCA cohort studied so far, we sought to provide foundational insights into GCA pathogenesis and identify potential therapeutic targets to address the significant clinical burden of this condition.

## Methods

### Study cohort

Temporal artery biopsies (TABs) from 69 patients with GCA and 10 controls were collected in hospitals from Italy, France, and Spain. All patients with GCA had a positive TAB, with transmural inflammation, and fulfilled the 1990 American College of Rheumatology classification criteria for this disease [13]. Clinical criteria for both inclusion or exclusion from the study is listed in *Supp Mat 1*. Control samples were obtained from GCA-suspected individuals whose TABs were negative. Moreover, the diagnosis of GCA was discarded based on additional clinical criteria and follow-up of at least one year. The main demographic and clinical characteristics of this cohort are described in *Table 1*. All participants signed an informed consent form in accordance with the ethical guidelines of the 1975 declaration of Helsinki. The protocol adhered to all ethical regulations and obtained approval from the ethics committee “Area Vasta Emilia Nord” of the Principal Investigator (protocol 777/2018/OSS/AUSLRE, date 10th October 2018) and from those of the participating institutions involved in this study.

**Table 1.** Clinical and demographic characteristics of the study cohort.

|  | Total | Cases | Controls |
| --- | --- | --- | --- |
| Cohort size (n) | 79 | 69 | 10 |
| Age (mean ± s.d.) | 75.54 ± 7.04 | 74.87 ± 6.72 | 80.20 ± 7.84 |
| Sex (Females) | 62 (70%) | 58 (84%) | 6 (60%) |
| GC treatment at TAB* | 43 (53%) | 41 (61%) | 2 (20%) |
| Treatment during follow-up period (GC / GC+TCZ) | - | 64 / 5 | - |
| Responders to treatment* | - | 32 (48%) | - |
| Any cranial symptoms*,<sup>1</sup> | 69 (90%) | 61 (91%) | 8 (80%) |
| Any visual symptoms*,<sup>2</sup> | 25 (32%) | 22 (33%) | 3 (30%) |
| Any systemic symptoms*,<sup>3</sup> | 38 (49%) | 38 (57%) | 0 (0%) |
| Polymyalgia rheumatica* | 30 (48%) | 27 (40%) | 3 (30%) |

Control samples were obtained from GCA-suspected individuals whose TABs were negative, in addition to further clinical criteria and follow-up of at least one year.
\*Information about clinical manifestations and response to treatment was not available for two patients with GCA
<sup>1</sup> Any cranial symptoms of the following: headache, scalp disesthesias, jaw claudication
<sup>2</sup> Any systemic symptoms of the following: fever, fatigue, weight loss ≥ 4Kg
<sup>3</sup> Any visual symptoms of the following: permanent visual loss, amaurosis fugax, diplopia
s.d., standard deviation; GC, glucocorticoids; TAB, temporal artery biopsy; TCZ, tocilizumab.

### Patients and public involvement

One Italian and two Spanish associations of patients were involved as patient research partners (PRPs): Association of patients of the Emilia Romagna (AMRER, Bologna, Italy), Spanish Forum of chronic patients (FEP, Vitoria, Spain) and Spanish Rheumatology League (LIRE, Madrid, Spain). They raised awareness about the disease in the respective associations, supervised the production of materials for patients in the context of the research, supervised the patient consent forms and translations, prepared leaflets about the disease and the research project and disseminated information about the disease and the project in the association social media.

### DNA methylation profiling

To preserve the *in situ* cellular and molecular composition, TABs samples were snap-frozen in liquid nitrogen and stored at –80°C within 30 minutes from the withdrawal. DNA from TABs was isolated with the DNA/RNA/Protein Purification Plus Kit (Norgen) following manufacturer’s instructions, and quantified with Qubit Fluorometric Quantification (ThermoFisher). DNA methylation profiling was performed using 500 ng of bisulfite-converted DNA hybridized to the Infinium MethylationEPIC BeadChip array (Illumina, Inc., San Diego, CA, USA) in accordance with the manufacturer’s protocol. This array, designed to target 850,000 CpG sites, provides coverage of 99% of annotated RefSeq genes. Probe fluorescence was measured using a BeadArray Reader (Illumina, USA).

### Quality controls and normalization

DNA methylation raw data was processed using *minfi* R package [14]. First, we checked for samples showing poor bisulfite conversion (detection p-value > 0.05) and those with discrepancies between recorded sex information and sex inferred from the data. Next, probes with a detection p-value inferior to 0.01 were removed. We also filtered out CpGs from chromosomes X and Y, as well as those probes containing either a SNP at the CpG interrogation site or at the single base extension site. Finally, cross-reactive probes reported in literature were also removed [15]. Probes were annotated using *IlluminaHumanMethylationEPICanno* (ver 5). Subsequently, we normalized raw methylation values using the stratified quantile normalization method.

Methylation levels were measured as β and M values. β values, representing the fraction of methylated probes on a scale from 0 to 1, were used for visual representation and biological interpretation. For statistical analysis, M values (log2-transformed β values) were calculated to achieve a normal distribution. To evaluate potential confounding effects, principal component analysis (PCA) was conducted (*Supp Mat 2*). In addition, PCA was also used to estimate potential outliers in the cohort. Samples exceeding a threshold of 4 standard deviations from the cluster centroids were excluded from further analyses.

### Genome-wide differential methylation analysis

We compared methylation levels of patients with GCA versus controls of all CpGs that passed quality controls using a eBayes moderated t-test from *limma R package* [16]. For this comparison, we conducted independent sensitivity tests to examine the contribution of all potential confounders (sex, age, treatment at TAB collection, sample plate, slide and well) to the analysis. Pearson correlation or Wilcoxon signed-rank test was applied depending on whether the variable of interest was continuous or categorical. Variables showing an adjusted p-value < 0.05 were considered to significantly contribute to DNA methylation profiles and were therefore included as covariates in the model (*Supp Table 1)*. For multiple-testing correction we applied a false discovery rate (FDR) of 0.05. The differential of β values (Δβ) between the groups considered in each comparison were estimated as the difference in the median values. Those CpGs sites with a adjusted p-value < 0.05 and |Δβ|>0.3 were considered as differentially methylated positions (DMPs). The enrichment of DMPs within genomic locations was analyzed using *EWAS toolkit* [17]. Furthermore, we assessed the presence of differentially methylated regions (DMRs), containing at least 5 significant CpGs, by using *DMRcate* [18].

### Pathway enrichment analysis

To explore the potential pathological relevance of the gene sets identified in our analyses, we conducted pathway enrichment analysis using the *EnrichR* tool [19]. The analysis was performed by querying two comprehensive databases: *GO Biological Process* (2023) [20] and *Bioplanet* (2019) [21]. Pathways needed to be enriched in at least 5 genes to be considered, and those presenting an adjusted p-value inferior to 0.05 were considered significant. As methylation changes can either upregulate or downregulate genes depending on their genomic context, we included all annotated genes in the pathway enrichment analysis, regardless of whether they contained hyper- or hypomethylated CpGs.

### Comprehensive overview of GCA-affected genes

In order to identify genes exhibiting transversal alterations across gene regulation, gene expression, and protein levels, we compared our results on DMPs and DMRs with reported findings from previous bulk omics studies in GCA. Specifically, we evaluated overlaps with: 1) DMPs previously reported in temporal arteries [12], 2) differentially expressed genes (DEGs) identified in temporal arteries [22], and 3) altered protein levels observed in plasma serum [23]. The summarized characteristics of the experimental design carried out in each of those studies are summarized in *Supp Table 2*.

## Results

### Specific epigenomic signature in GCA-affected arteries

After quality controls, 770,921 CpGs were analyzed and all samples were included in the analysis. Notably, 89.2% of CpGs showing significantly altered methylation level in GCA-affected arteries in a previous study [12] were also significant in our results (p_adj_ < 0.05) and showed a strong correlation in their Δβ values between studies (Spearman correlation=0.75, p < 2.2×10^-16^). In addition, all overlapping significant CpGs displayed methylation changes in the same direction (*Supp Mat 3*). Due to the increased statistical power in our experimental design, a total of 228,152 CpG sites presented significant changes in methylation (p_adj_ < 0.05), demonstrating an extensive epigenetic alteration present in the GCA-affected artery. Therefore, we focused on those significant CpGs showing major changes in methylation (|Δβ|>0.3, p_adj_<0.05), comprising 3,644 DMPs, where 81.1% of them represent novel DMPs for arteries affected by GCA. Among these, 2,680 were hypermethylated, and 964 were hypomethylated (*Fig 1, Supp Table 3*). These DMPs were predominantly located in OpenSea regions and gene bodies (*Supp Table 4*), and were annotated to 1,517 different genes, including relevant genes involved in GCA pathological mechanisms related to inflammatory response and vascular remodeling, such as *IL6R, IL2RA, STAT3* and *JAG1*. Pathway analysis of these DMPs showed 78 significant pathways (*Supp Table 5-6*), including fundamental immune pathways related to interleukin (IL) 2 and CTLA4 signaling, as well as mechanisms linked to the angiogenic process, such as integrins affecting angiogenesis and CXCR4 signaling (*Fig 2*).

**Figure 1.**
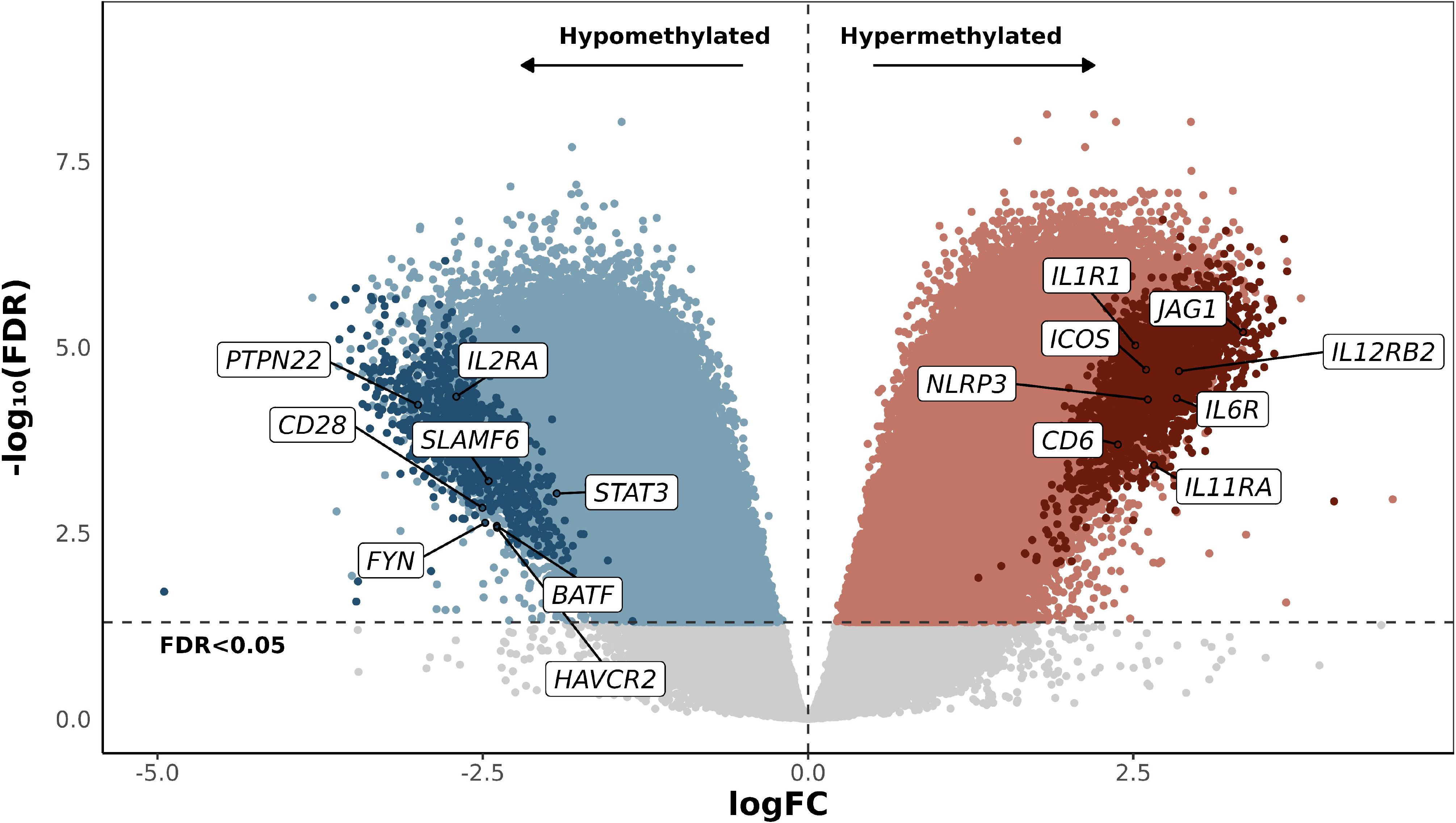
Volcano plot of the epigenome-wide comparison between patients with GCA and controls. Gray points represent the non-significant (adjusted p > 0.05) CpGs, blue and red points represent significant hypomethylated and hypermethylated positions, respectively. Darker-colored points represent differentially methylated positions (DMPs), significant CpGs showing |Δβ|>0.3. FC, fold change; FDR, false discovery rate.

**Figure 2.**
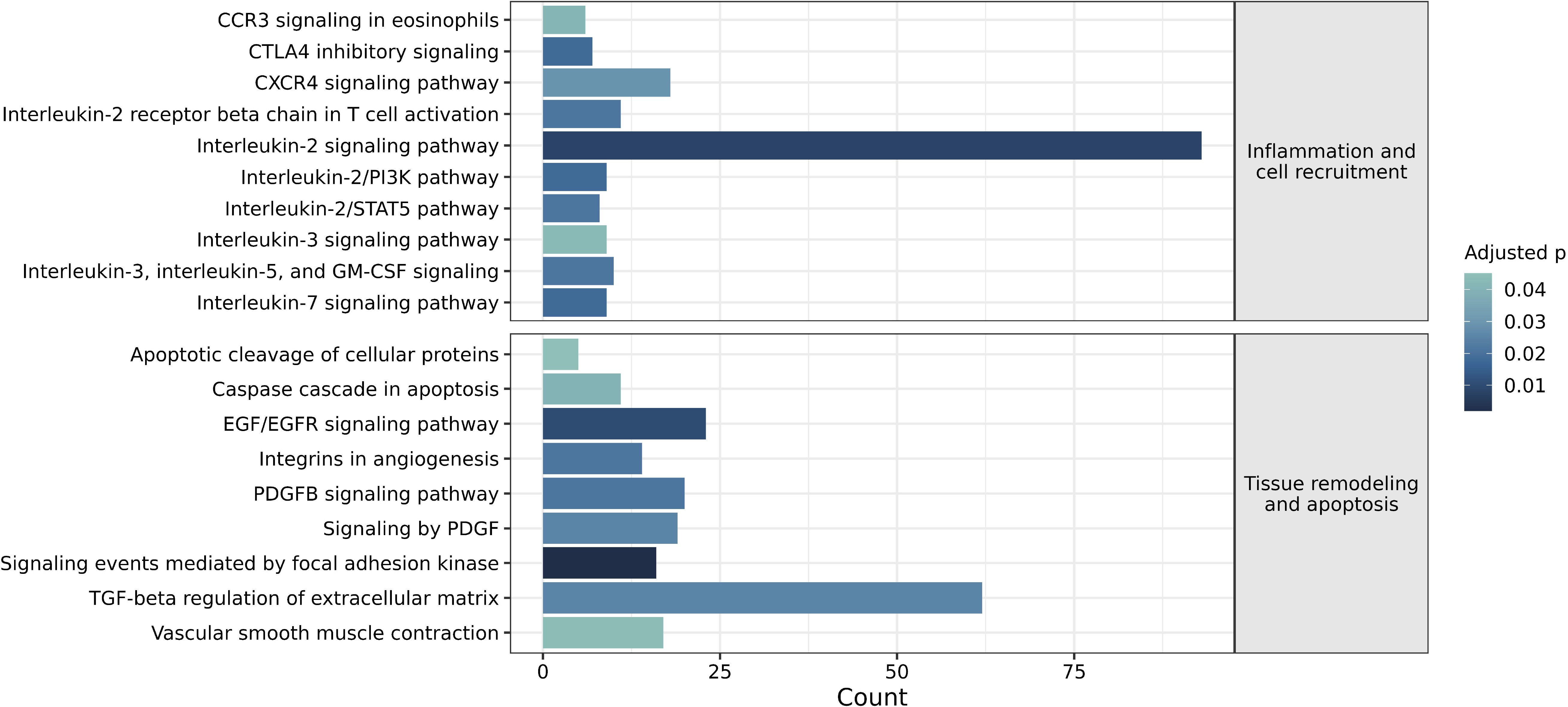
Highlighted significant pathways (adjusted p < 0.05) of the 1,517 genes annotated from differentially methylated positions.

### Epigenetic altered regions and affected pathways in GCA

To determine consistent methylation changes in the epigenome, we conducted a DMR analysis, identifying 12,899 significant regions associated with GCA (p_FDR_<0.05). Among these, 5,393 were hypomethylated and 7,506 hypermethylated, with a mean size of 9.53 CpGs per region. A subset of 918 DMRs showed substantial methylation changes (|mean Δβ| > 0.2) and were annotated to 935 genes (*Supp Table 7*), comprising 282 additional genes from those identified by annotation of DMPs. This gene set included critical immune and GCA-associated genes, such as *IL17B, IL11RA, HLA-DPA1, TNF*, and *PDGFA*, further supporting a strong alteration of immune activation and tissue remodeling. Pathway enrichment analysis highlighted disruptions in key arterial pathways (155 significant pathways; *Supp Table 8-9*). Notably, pathways related to arterial remodeling were enriched, including regulation of growth factors VEGF and PDGF. Additionally, signaling pathways relevant for arterial cell migration and immune cell activation were affected, such as leukocyte transendothelial migration, regulation of T cell activation and natural killer cell-mediated cytotoxicity (*Fig 3*). These findings emphasize the extensive epigenetic reprogramming that drives the local arterial response in the context of GCA pathogenesis.

**Figure 3.**
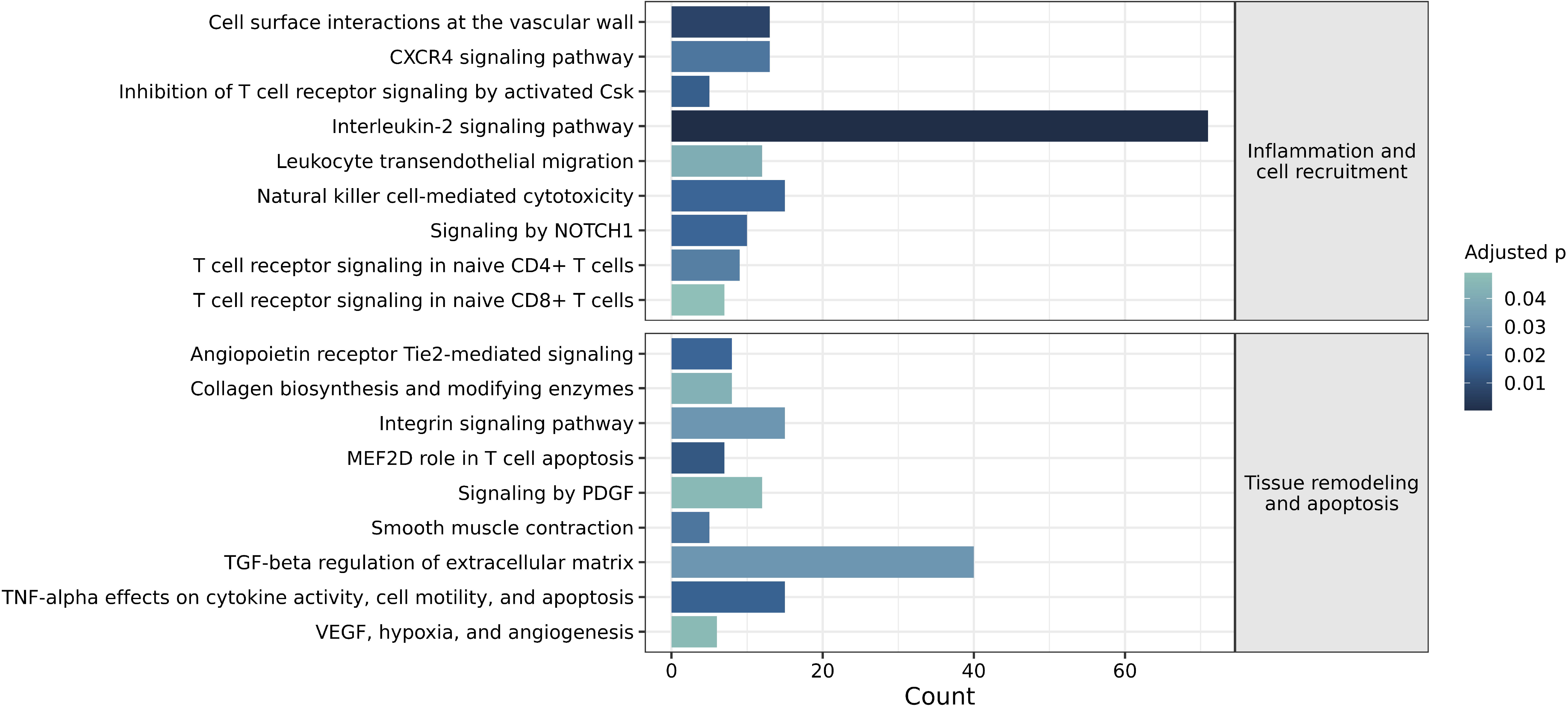
Highlighted significant pathways (adjusted p < 0.05) of the 935 genes annotated from differentially methylated regions.

### Consistently altered genes in GCA omics studies

To contextualize our findings, we evaluated the affected genes identified in this study against previously reported omics datasets on GCA. Notably, we identified 37 genes consistently altered in GCA-affected arteries across three major epigenomics and transcriptomics studies of this tissue, including this current work (*Fig 4*). This list comprises genes annotated either by DMPs or DMRs from our results, DMPs reported by Coit et al. [12], and genes classified as differentially expressed in Ferrigno et al [22]. These 37 genes carried a robust body of omics evidence supporting their involvement in the pathophysiology of GCA, and contain genes related to immune signaling (predominantly related to T cell mechanisms), tyrosine kinases, transcription factors, integrins, as well as other elements. While the list includes several genes known to be affected in GCA pathogenesis, such as *CD28, IL2RA*, and *TNF*, it also highlights many genes whose potential role in the pathogenesis of this disease have not been explored, such as *ICOS, GZMA, IRF5, TLR1* and *FYN*, among others. *BTLA* and *SLAMF1* were particularly outstanding, as they also exhibited significantly altered protein levels in plasma serum from patients with GCA [23]. These findings warrant further investigation given their consistent alterations across independent GCA omics studies.

**Figure 4.**
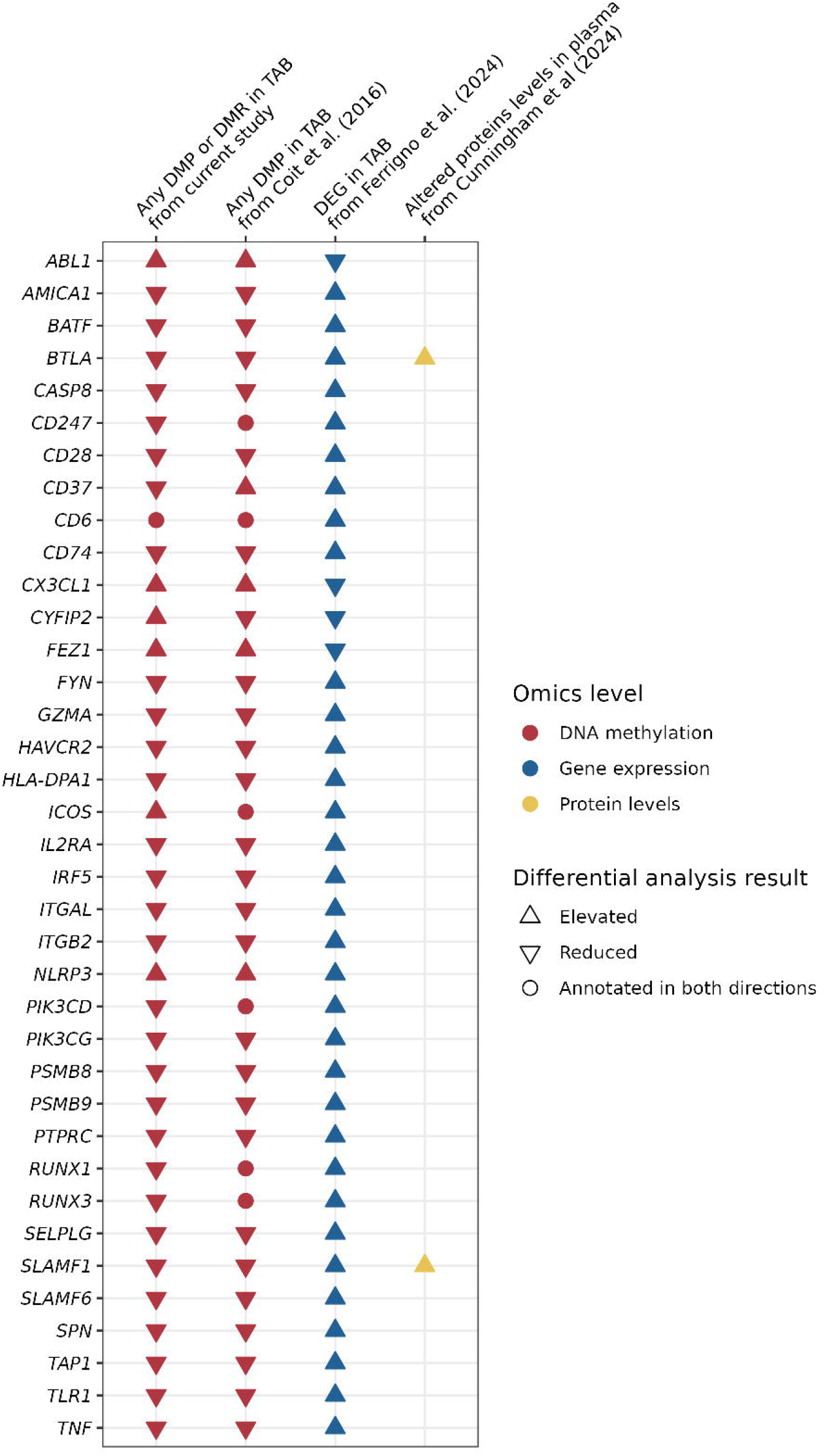
Reported alterations of gene regulation, gene expression and/or protein abundance levels in independent GCA omics studies for the 37 consistently reported genes in GCA-affected arteries. DMP, differentially methylated position; DMR, differentially methylated region; DEG, differentially expressed gene.

## Discussion

This study represents the largest genome-wide DNA methylation study of GCA-affected arteries conducted to date, providing comprehensive insights into the severe epigenetic dysregulation within arterial tissue underlying this pathology. We analyzed the most notable alterations in the GCA-associated signature, revealing major changes in critical immune and vascular pathways, especially those related to immune activation and vascular remodeling. Furthermore, by leveraging findings from epigenomics and transcriptomics studies, we identified key immune-related genes found consistently altered in GCA-affected arteries, comprising 37 genes that may play a fundamental role in driving this condition.

Interpreting omics data in a complex condition such as GCA is challenging due to the sheer number of genes implicated. To address this, we prioritized genes with strong replicability across independent omics studies using TAB samples. Our integrative approach yielded a shortlist of 37 genes exhibiting significant methylation changes (replicated in two independent studies) and altered gene expression levels. This shortlist encompasses a robust set of genes potentially central to GCA, being many of them not previously proposed to play a specific role in the pathogenesis of this disease. Among these, we observed a strong representation of co-stimulatory proteins involved in T cell and antigen-presenting cell interactions, including CD28, CD247, BTLA, ICOS, and SLAM proteins (SLAMF1, SLAMF6). Additionally, immune-related signaling molecules such as TNF, FYN, ABL1, PIK3CD, and PIK3CG, as well as transcription factors including RUNX1, RUNX3, and BATF, were prominent. One particularly noteworthy gene in this set is *NLRP3*, which encodes an intracellular pattern recognition receptor that detects pathogens, endogenous damage, and environmental irritants. After activation, it promotes the formation of the NLRP3 inflammasome that leads to proinflammatory signaling and apoptosis. This innate immunity-related pathway plays a central role in cardiovascular diseases [24] and is gaining recognition for its role in autoimmune diseases [25], therefore its potential contribution to GCA pathogenesis deserves further study, especially given its established value as a therapeutic target for conditions presenting chronic inflammation [26].

T cell exhaustion is a process in which T cells, following chronic exposure to antigens and persistent inflammatory signals, progressively lose their effector functions and cytotoxicity while upregulating inhibitory receptors [27]. While this phenotype is traditionally linked to chronic infections and cancer, it is now increasingly recognized in autoimmune and inflammatory diseases [27]. Our study, along with other GCA omics studies, identified several markers of exhausted T cells as being altered in GCA-affected arteries at both gene regulation and expression levels. Notable examples include *SLAMF6, HAVCR2* (also known as TIM-3), and *CX3CL1* (the ligand for CX3CR1), specific markers of the exhausted T cell phenotype [28]. Additionally, we also observed consistently reported alterations in *BTLA* and *BATF*, regulators of exhausted T cells differentiation [29]. It is also important to note that PD-1 and CTLA4, immune checkpoint molecules implicated in regulating inflammation, are central to the progression of GCA [4, 30] and also deeply linked to the development of exhausted T cells [27]. Furthermore, the CXCR4 pathway, significant in our pathway enrichment results, is also related to exhausted T cell differentiation, as it has been reported that CXCR4 can promote T cell exhaustion via JAK2/STAT3 [31]. Based on the convergence of GCA-related pathways and T cell differentiation, as well as the replicability of reported alteration of specific markers of exhausted T cells across GCA studies, we hypothesize that the microenvironment of GCA-affected arteries—characterized by persistent inflammation, hypoxia, and extracellular matrix damage—provides favorable conditions for the emergence and maintenance of exhausted T cell phenotypes that would impede the ability of the immune system to regulate the local inflammation occurring in the affected tissue.

While T cell exhaustion has been primarily described in CD8+ T cells, the role of this T cell subtype in GCA remains poorly understood despite evidence of their presence in inflamed arterial tissue [32]. Emerging evidence suggests that CD4+ T cells can also exhibit an exhausted phenotype [33], and given the established importance of CD4+ T cells in GCA pathogenesis, it would be particularly relevant to investigate whether exhausted subsets of both CD8+ and CD4+ T cells contribute to disease progression. Understanding the functional implications of T cell exhaustion in GCA could reveal novel mechanisms of immune dysregulation and provide opportunities for targeted therapeutic interventions aimed at restoring T cell function or mitigating exhaustion pathways, a promising therapeutic strategy currently under active investigation in other conditions [34, 35].

Regarding epigenetic alterations newly identified in this current study, pathway analysis revealed a broad spectrum of disrupted mechanisms in the context of GCA. Among these, the IL-2 signaling pathway emerged prominently, with the highest number of affected genes (n=93) being annotated to this pathway. This finding emphasizes the contribution of IL-2 signaling in GCA pathophysiology, associated with vascular persistent Th1 cell signature [36]. Notably, low-dose IL-2 therapy has shown significant promise in helping regulate inflammatory responses across a broad range of IMIDs [37]. These results suggest GCA as a compelling candidate for exploring the therapeutic potential of IL-2-based treatments. Similarly, the IL-7 signaling pathway, also related to IL-2, was found to be significantly enriched in our results. While IL-7 is well-documented to play a role in other IMIDs [38] and is known to affect immune cell populations, e.g. promoting T-cell apoptosis [39], its involvement in GCA pathogenesis represents a novel finding showing specific therapeutic potential [40]. Furthermore, our study also found a significant change in the regulation of *IL11RA*, which supports the recently proposed contribution of IL-11 in GCA pathogenesis by a multi-omics study of blood-derived GCA monocytes [8]. Finally, another notable finding was the novel association of the CXCR4 signaling pathway with this disease, an essential regulator of local inflammation [41]. CXCR4 is instrumental in preserving arterial integrity, regulating vascular inflammation, promoting angiogenesis and the recruitment of inflammatory cells [42, 43], processes that are central to the pathophysiology of GCA. The implication of CXCR4 in GCA pathogenesis also shows great therapeutic promise, as CXCR4-targeted therapies are under active research [44]. These findings reinforce the relevance of local immune and vascular signaling in GCA and suggest that targeting pathways such as CXCR4 and IL-2 could be pertinent for therapeutic innovation in this disease.

Our study offers valuable insights into the epigenetic changes in GCA-affected arteries, leveraging well-powered genome-wide DNA methylation profiling and multi-omics comparison to provide a comprehensive overview of this disease. The use of a notable cohort and robust analytical methods allowed us to identify novel pathways and mechanisms, such as T cell exhaustion and CXCR4 signaling. However, several limitations should be acknowledged to fully contextualize these findings The omics evidence presented here provides statistical evidence but requires further functional studies to clarify how the proposed pathways and genes mechanistically contribute to GCA pathogenesis. Moreover, the substantial clinical heterogeneity of patients with GCA posed challenges for our analysis. Although we identified a robust epigenetic signature associated with GCA, limitations in the sample size of our cohort prevented the possibility of studying the effect of particular clinical manifestations in DNA methylation, restricting our ability to detect epigenetic patterns linked to particular phenotypes. These limitations highlight the importance of future research involving larger, clinically well-characterized cohorts and experimental validation to enhance our understanding of the molecular mechanisms driving GCA pathogenesis.

This study represents a methodologically robust analysis into the epigenetic landscape of GCA-affected arteries, leveraging genome-wide DNA methylation profiling and previous omics findings to provide novel insights into the pathogenesis of this disease. Among the key results, the potential involvement of exhausted T cells in GCA pathophysiology represents an intriguing discovery, representing a new potential agent of the immune dysfunction present in GCA-affected arterial tissue. These findings enhance our understanding of GCA pathogenesis and highlight therapeutic targets, paving the way for improved treatment strategies.

## Acknowledgements

Acknowledgements

We express our gratitude to Sofia Vargas for her exceptional technical support, as well as to all the patients and control donors for their indispensable cooperation. This study was presented at the EULAR 2024 Congress under the title “OP0025 Genome-wide DNA methylation study reveals specific signatures in the arterial tissue of giant cell arteritis patients” (DOI: 10.1136/annrheumdis-2024-eular.4859). This research is part of the doctoral degree awarded to GB-Y, within the Biomedicine program from the University of Granada.

## Data availability statement

Summary results will be made available upon reasonable request. Individual-level data cannot be shared due to privacy and ethical considerations.

## Contributors

GB-Y, AM and LO-F: data analysis, interpretation of the results and manuscript drafting. EE-M and LCT-C: data analysis. MAG-G, SCa, GG, DS, PL, SF, MB, AR, FM: sample and data collection. CS, NP and SCr: study design, sample and data collection. JM: study design, interpretation of the results and manuscript drafting. JM, SCr and LO-F: guarantors of the study. All authors read and approved the manuscript.

## Patient consent for publication

Not applicable.

## Ethics approval

The study was approved by the ethical committees of all institutions involved in this study. Participants gave informed consent to participate in the study before taking part.

