## Supplementary material for "Genome-wide DNA methylation study reveals specific signatures in the affected arterial tissue of giant cell arteritis patients": Supp materials

SUPPLEMENTARY MATERIALS

Genome-wide DNA methylation study reveals specific signatures in the affected arterial tissue of giant cell arteritis patients	0

1. Additional exclusion and inclusion clinical criteria	2

2. PCA distribution plots before and after normalization	3

3. Sensitivity test results for differential methylation analyses	4

### 1. Additional exclusion and inclusion clinical criteria

- **Inclusion criteria**

Patients had to meet the following criteria for study entry:

1. Diagnosis of GCA confirmed by TABs showing transmural inflammation.
2. Age >= 50 years.
3. GC-therapy naïve or previous treatment with GCs for a maximum of 14 days at a maximal dose of 60 mg/day of prednisone or equivalent at the time of TABs.
4. Able and willing to provide written informed consent and to comply with the study protocol according to the physicians’ judgment.

- **Exclusion criteria**

Patients were excluded for the study if they meet any of the following criteria:

1. Use of GCs for more than 14 days by the time of TABs.
2. Intravenous methylprednisolone pulses ≤ 3 months before TABs.
3. Active inflammatory rheumatic diseases other than GCA or polymyalgia rheumatica.
4. Previous treatment with TCZ or other biologic agents.
5. Treatment with hydroxychloroquine, cyclosporine A, azathioprine, mycophenolate mofetil, methotrexate, cyclophosphamide within 4 weeks of TABs.
6. Evidence of significant and/or uncontrolled concomitant conditions which, in the investigator’s opinion, would preclude patients participation or negatively impact the benefit-risk ratio including incoming major surgery.

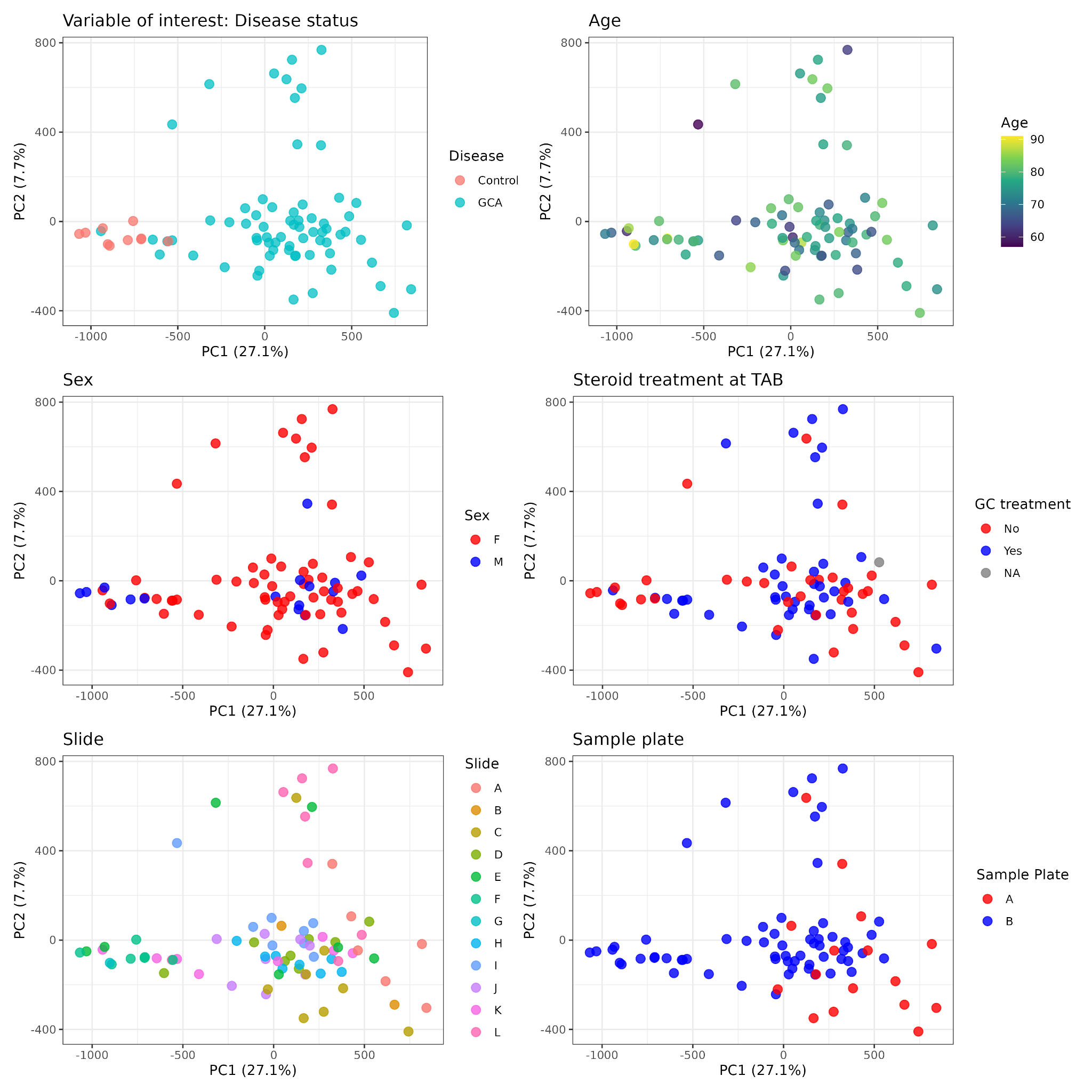

### 2. PCA distribution after normalization

**Supplementary Material 2.** PCA cohort distribution according, in order from left to right and top to bottom, to: disease status, age, sex, glucocorticoid treatment at the moment of the temporal artery biopsy, methylation sample slide and methylation sample plate.

### 3. Effect size correlation between GCA DNA methylation studies

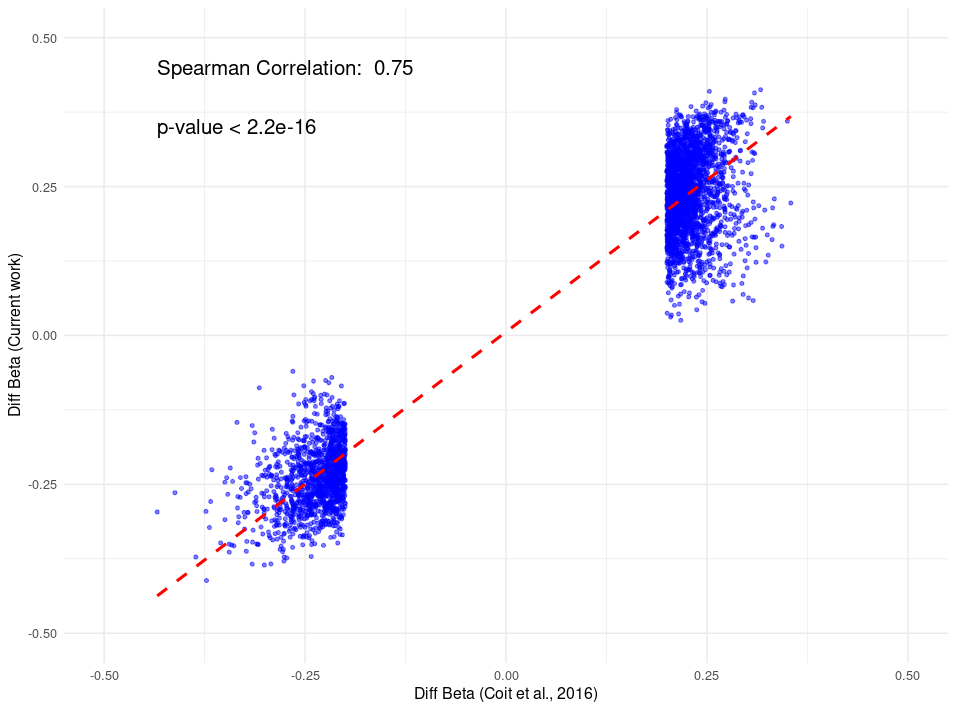

Spearman correlation between the Δβ values of the differentially-methylated positions in GCA-affected arteries reported in a previous study*, and their Δβ values calculated in the current study, for those DMPs that were also present in the current work (89,2%).

**Coit P, De Lott LB, Nan B, Elner VM, Sawalha AH. DNA methylation analysis of the temporal artery microenvironment in giant cell arteritis. Ann Rheum Dis. 2016 Jun;75(6):1196-202.*
